# Influence of E/I balance and pruning in peri-personal space differences in schizophrenia: a computational approach

**DOI:** 10.1101/2020.11.06.371450

**Authors:** Renato Paredes, Francesca Ferri, Peggy Seriès

**Affiliations:** The University of Edinburgh, School of Informatics, 10 Crichton Street, Edinburgh, United Kingdom; Department of Neuroscience, Imaging and Clinical Sciences, University of Chieti-Pescara, Chieti, Italy

**Keywords:** peri-personal space, neural network, E/I balance, computational psychiatry

## Abstract

The encoding of the space close to the body, named peri-personal space (PPS), is thought to play a crucial role in the unusual experiences of the self observed in schizophrenia (SCZ). However, it is unclear why SCZ patients and high schizotypal (H-SPQ) individuals present a narrower PPS and why the boundaries of the PPS are more sharply defined in patients. We hypothesise that the unusual PPS representation observed in SCZ is caused by an imbalance of excitation and inhibition (E/I) in recurrent synapses of unisensory neurons or an impairment of bottom-up and top-down connectivity between unisensory and multisensory neurons. These hypotheses were tested computationally by manipulating the effects of E/I imbalance, feedback weights and synaptic density in the network. Using simulations we explored the effects of such impairments in the PPS representation generated by the network and fitted the model to behavioural data. We found that increased excitation of sensory neurons could account for the smaller PPS observed in SCZ and H-SPQ, whereas a decrease of synaptic density caused the sharp definition of the PPS observed in SCZ. We propose a novel conceptual model of PPS representation in the SCZ spectrum that can account for alterations in self-world demarcation, failures in tactile discrimination and symptoms observed in patients.

## 1. Introduction

The brain encodes the space that surrounds the body to be able to interact with the environment. These space representations are split into regions according to their distance from the body: peri-personal space (PPS) (i.e. the space reachable by hand) and extrapersonal space (EPS) (i.e. the space that cannot be reached by hand) (van der Stoep et al., 2016). This classification is supported by evidence of fronto-parietal neurons of macaques (Fogassi et al., 1996; Graziano et al., 1999) and humans (Bernasconi et al., 2018; Brozzoli et al., 2014; Ferri et al., 2015; Gentile et al., 2013; Làdavas et al., 1998; Noel et al., 2018c; di Pellegrino et al., 1997) responding stronger to visual and auditory stimulation that occurs near the body (see Grivaz et al. (2017) for an extensive review).

Maintaining a representation of the PPS is thought to allow animals to respond faster to stimuli near the body as a mechanism against threats from the environment (Graziano and Cooke, 2006). Evidence from bodily self-consciousness experiments ^1^(e.g. Enfacement Illusion, Body-Swap Illusion and Full Body Illusion) have led to the hypothesis that PPS representation is relevant to bodily self-consciousness processes in the brain (Blanke et al., 2015; Noel et al., 2018a). More recently, it has been proposed that PPS representation is crucial in the causal inference process involved in bodily self-consciousness because it couples body-related information and surrounding exteroceptive signals (Noel et al., 2018b).

Patients with schizophrenia are thought to hold a weaker or more variable PPS representation (Noel et al., 2017). This view is based on the observation that patients with schizophrenia are more prone to experience bodily self-aberrations such as the Rubber Hand Illusion (RHI) ^2^ (Thakkar et al., 2011) and the Pinocchio Illusion (PI) ^3^ (Michael and Park, 2016). As a consequence, an initial working model was proposed suggesting that space representations in schizophrenia (SCZ) are characterised by a shallow gradient dividing the PPS and the extrapersonal space (Noel et al., 2017). A shallower demarcation of the PPS is thought be an indicator of reduced self demarcation (i.e. confusion of boundaries between self and others and permeability of self-world boundaries).

Recently, this working model was directly tested and experiments revealed that patients with schizophrenia and participants scoring high for schizotypy as measured by the SPQ questionnaire (H-SPQ) present a narrower PPS compared to healthy controls and low-schizotypy individuals (Di Cosmo et al., 2017). However, contrary to conceptual predictions, it was found that patients with schizophrenia present sharper PPS boundaries (i.e. steeper gradients dividing the PPS and the extrapersonal space) compared to controls.

### 1.1. Problem statement

The neural basis underlying these differences observed in patients with schizophrenia and individuals with high schizotypal traits is unknown. There has not been an in depth examination of the disagreement between the initial working model of PPS representation in SCZ (Noel et al., 2017) and the aforementioned experimental evidence (Di Cosmo et al., 2017). Particularly, a comprehensive explanation of why patients with schizophrenia and individuals with schizotypal traits present a narrower PPS is lacking. Similarly, there is as yet no theoretical understanding of why PPS boundaries are sharply defined in patients with schizophrenia.

Similarly, the mechanisms behind the propensity of patients to manifest bodily self-aberrations are unclear. Although the literature points out a relationship between tactile sensitivity, self-disturbances and failures in bodily self-consciousness (Chang and Lenzenweger, 2001, 2005; Costantini et al., 2020; Ferri et al., 2016; Michael and Park, 2016), it is unknown how this is related to the encoding of the PPS and the unusual PPS representation found in patients with schizophrenia and individuals with high-schizotypal traits. Moreover, there is not a clear understanding of how a narrow PPS is related to symptoms of schizophrenia and schizotypal traits.

### 1.2. Objectives and Hypothesis

The current study aims at implementing a computational model that can account for PPS representation in SCZ and H-SPQ (Di Cosmo et al., 2017) and is compatible with state of the art computational modelling of psychosis (Friston et al., 2016; Sterzer et al., 2018). For this purpose, a recurrent neural network model of PPS (Magosso et al., 2010; Serino et al., 2015a) is adapted to reproduce PPS changes observed in SCZ (i.e. size reduction and boundaries sharpening) in a simulated audio-tactile behavioural task. The simulated results are compared against experimental data of patients with schizophrenia and individuals with high schizotypal traits (Di Cosmo et al., 2017). The adapted models are then used to generate predictions regarding tactile discrimination and bodily self-aberrations.

We hypothesise that a narrower PPS representation in SCZ spectrum disorders is a result of an E/I imbalance in recurrent synapses among unisensory neurons, in agreement with neurobiological evidence (Jardri et al., 2016). Moreover, in line with anatomical observations, we expect that PPS boundaries sharpening in SCZ may be due to an impairment of bottom-up and top-down connectivity between unisensory and multisensory neurons (Ellison-Wright and Bullmore, 2009; Harrison, 1999). Finally, we conjecture that both E/I modulation and connectivity impairments are related to deficits in tactile discrimination and bodily self-aberrations observed in patients.

## 2. Methods

### 2.1. Experimental task

The computational model simulates the experiment conducted on patients with schizophrenia and H-SPQ individuals in the aforementioned study (Di Cosmo et al., 2017) (see Figure 1 for an illustration). This experiment consisted of presenting approaching sounds (i.e. 3000 ms sounds with exponentially increasing intensity from 55 to 70 dB) to participants combined with a tactile stimulation on the right hand administered at different delays from the sound onset, ranging from 300 ms to 2700 ms (see Ferri et al. (2015) for an extensive description). The manipulation of the delay of the tactile stimulus produces the subjective perception that the longer the delay, the closer to the participant’s hand the sound is perceived.

**Figure 1:**
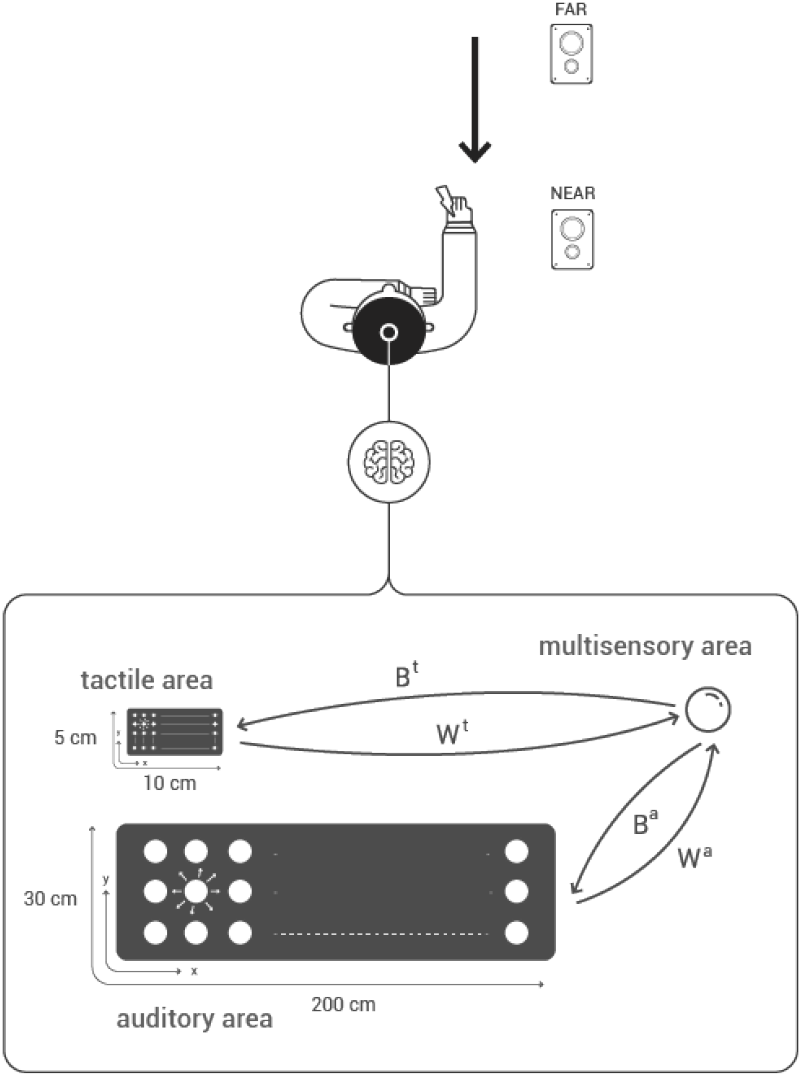
Audio-tactile experimental paradigm and model of PPS audio-tactile representation. In this experimental setup, two speakers are placed in front of the participant and an electrode is fit on his right hand. The black arrow indicates the subjective perception of sounds moving towards the participant respectively with tactile stimulation administered at different delays from the sound onset. At every trial, participants were required to respond as fast as possible to the tactile stimulation by pressing a button with their left index finger. Enhanced tactile RTs are observed when the sounds are perceived within the boundaries of the PPS. The network model is composed of two unisensory areas (tactile and auditory) connected with a multisensory area. The unisensory areas are arranged to encode the space of the hand (10 cm x 5 cm) and the external auditory space (200 cm x 30 cm) respectively.

At every trial, participants were required to respond as fast as possible to the tactile stimulation by pressing a button with their left index finger, while sounds were delivered, which participants were told were task-irrelevant. In such paradigm, faster responses to the tactile stimulation are typically observed when the sound is subjectively perceived in the space near the body. This is thought to be related to the activity of multimodal neurons in parieto-frontal networks projecting back to somatosensory regions of the cortex (Cléry et al., 2015; Graziano and Cooke, 2006; Grivaz et al., 2017).

For each participant, the average reaction times (RTs) as a function of the delay at which the tactile stimuli were administered were fitted by a sigmoidal function. This function can be interpreted as a representation of the individual PPS: fast responses correspond to sensory areas within the PPS while slow responses correspond to area outside the PPS. Hence, the central point (CP) of the curve is interpreted as the extent of the PPS, whereas the slope is understood as the steepness of the PPS boundary. In the empirical study (Di Cosmo et al., 2017), it was found that SCZ patients presented higher CPs (i.e. a smaller PPS) and steeper slopes than HC, whereas H-SPQ individuals displayed only higher CPs in comparison with L-SPQ individuals (see Figure 3).

**Figure 2:**
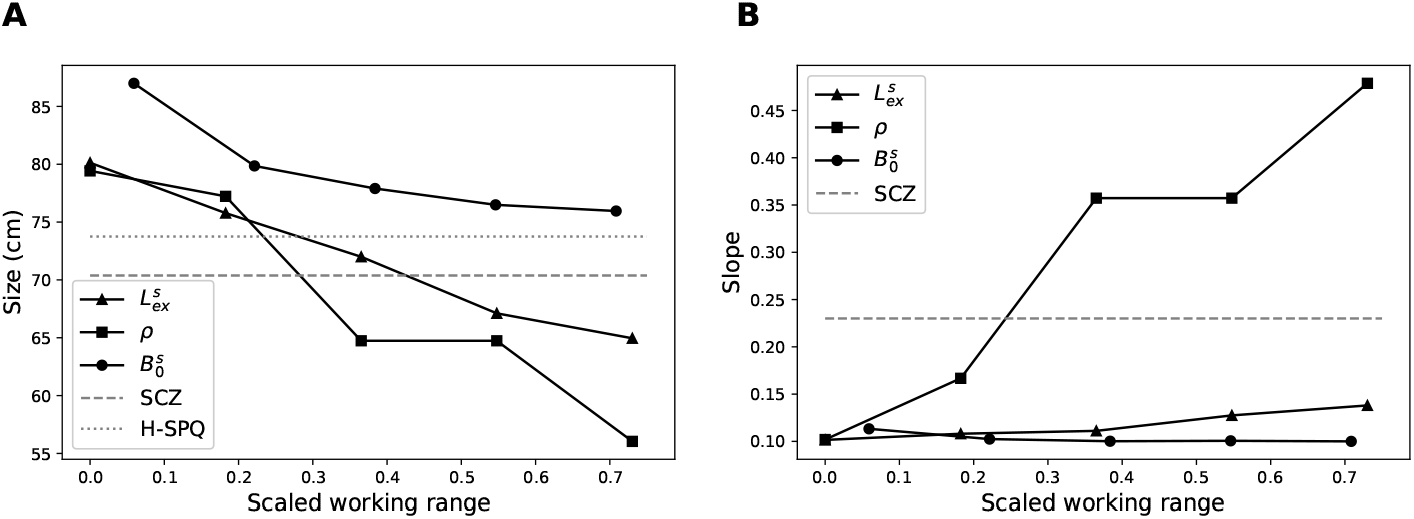
Effects of potential neural impairments in the size and slope of the PPS representation generated by the HC model. **Panels A** and **B** show the effect of systematic variation of parameters in the range at which they produce sigmoid-like PPS representations (these ranges were scaled to facilitate visual comparison). The dashed lines indicate the size and slope reported in the experimental study (Di Cosmo et al., 2017). The size reduction observed in SCZ and H-SPQ could be reproduced by either an increase in lateral excitation (*L_ex_*) or pruning of feedforward synapses (*ρ*). In contrast, the sharpening of the slope observed in SCZ could only be reproduced by the pruning mechanism.

**Figure 3:**
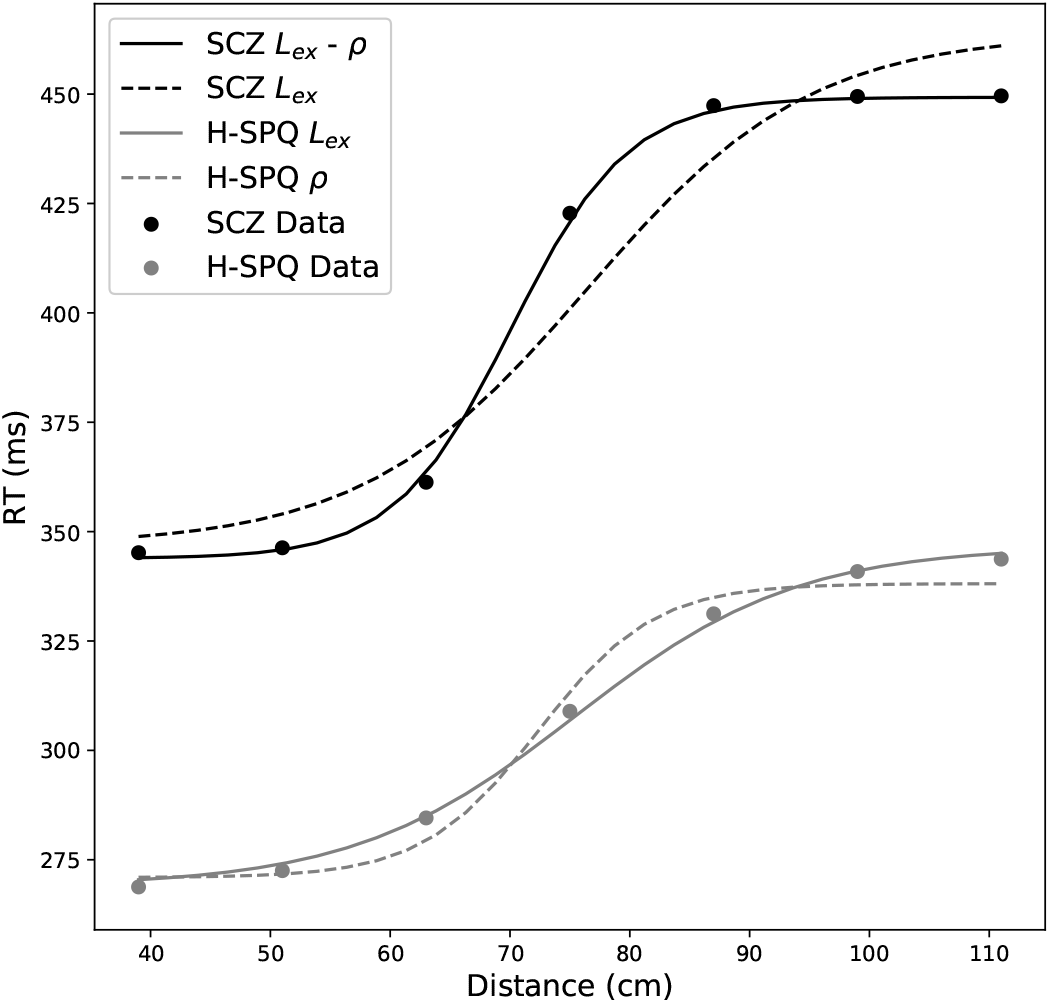
PPS representations generated by the SCZ and H-SPQ network models identified by the fitting procedure. The solid lines depict the sigmoid fit obtained out of the data generated by the models. The dots represent the discretisation of the sigmoid fit obtained out of the data collected in the experiment (Di Cosmo et al., 2017). The quantitative fitting of the HC model using only 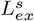 is sufficient to reproduce the H-SPQ data (grey). In contrast, both 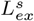 and *ρ* are required to generate a close match to the SCZ data (black).

### 2.2. Neural network model of PPS representation

We model the task by adapting the neural network model proposed by (Serino et al., 2015a). The main modifications were a reduction in the number of tactile neurons, the elimination of noise sources and the introduction of a mapping between neural activation and behavioural reaction times. Importantly, these three changes allowed to fit the model to behavioural data.

In short, the model describes two unisensory areas (auditory and tactile) connected with a multisensory area that holds multisensory representations of the PPS (see Figure 1 for an illustration). Neurons in both unisensory areas are characterised by their receptive field (RF) defined in hand-centred coordinates (along the horizontal and vertical axis) and respond to stimulation at specific spatial coordinates in relation to the hand. The tactile area is composed of 200 neurons (disposed in a *M^t^* = 20 x *N^t^* = 10 grid) that encode a skin portion of 10 cm x 5 cm corresponding to the left hand of an individual. The auditory area is composed of 60 neurons (disposed in a *M^a^* = 20 x *N^a^* = 3 grid) that cover an auditory space of 200 cm x 30 cm on and around the hand.

For simplicity, the multisensory area is composed of a single neuron connected to all neurons in both auditory and tactile areas, both via feedforward and feedback synapses. Hence, the RF of this multisensory neuron has a wider spatial extension compared to individual unisensory neurons. The multisensory neuron has stronger feedforward and feedback synapses from/to the auditory neurons encoding areas close to the modelled hand (i.e. the PPS). The multisensory neuron responses are consistent with observed responses of neurons located in parietal, temporo-parietal and premotor regions which respond to tactile stimuli in the body and to auditory stimuli presented close, but not far from the body (Bernasconi et al., 2018; Graziano et al., 1999; Grivaz et al., 2017; Schlack et al., 2005).

Using this model, RTs of healthy participants in the audio-tactile experimental paradigm described above (Ferri et al., 2015) can be simulated. The reaction time of the network was registered at each distance point as the time at which any neuron of the tactile area reached 90% of its maximum activation state. An extensive description of the model and values of the parameters can be found in the Supplementary Material ^4^.

### 2.3. Experiment simulation in the network

The experiment was simulated by holding the following assumptions: First, the apparent velocity of the sound was 30 cm/s ^5^. Second, the inputs to the network, representing the stimuli, were configured to last 100 ms for both auditory and tactile stimuli (mimicking possible responses of input neurons to the audio-tactile stimulation which is much shorter (i.e. 100 *μ*s)) (Serino et al., 2015a). Third, model simulations evaluated RTs for tactile stimuli presented with 7 different delays from the sound onset ranging from 300 ms to 2700 ms, which according to our first assumption correspond to 7 subjectively perceived sound distances between 39 cm and 111 cm from the hand. Fourth, those subjectively perceived sound distances were implemented as if the sound stimuli were physically presented at different distances from the modelled hand, as in Serino et al. (2015a).

Fifth, it was assumed that the network model does not capture the entire neural process that causes the motor responses (i.e. button pressing) examined in the task. As a consequence, the scale of the network’s RT does not match exactly with RT found in human participants. To re-scale the reactions times and make them comparable to human data, a linear regression is applied to the raw RTs of the network following the procedure described in (Bogacz and Cohen, 2004).

Sixth, the parameters that define the feedforward and feedback auditory synaptic weights outside the space near the hand were fitted to the average sigmoid fit obtained in the HC group (Di Cosmo et al., 2017) (see Supplementary materials for a full description of the fitting procedure). The purpose of this manipulation was to calibrate the model to reproduce PPS representations of healthy individuals that closely resemble the ones observed in the empirical study (RMSE = 0.37 ms). Hence, in the following this network setup will be denominated HC model and will be taken as the baseline.

### 2.4. Modelling the influence of SCZ and H-SPQ in the network

The HC model setup was considered as a starting point to introduce impairments such as those observed in SCZ and H-SPQ (Di Cosmo et al., 2017). More precisely, we aimed to determine what impairments cause the size reduction of the PPS and the sharpening of its boundary. For this purpose, we evaluated three consistently observed impairments in the SCZ spectrum: excitation/inhibition (E/I) imbalance (Jardri et al., 2016), failures in top-down signalling (Sterzer et al., 2018) and synaptic density decrease (Ellison-Wright and Bullmore, 2009).

#### 2.4.1. Modulation of the E/I balance

The modulation of the E/I balance was implemented by increasing or decreasing the strength of recurrent excitatory connectivity. This was done by modifying the value of the parameter 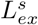, defined in equation 1:

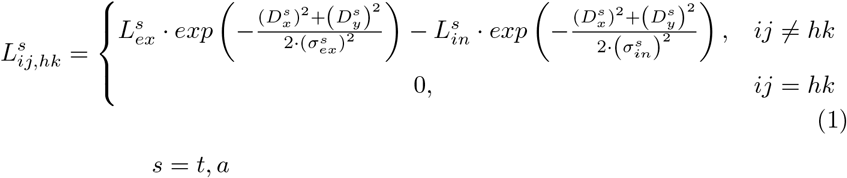

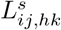 denotes the weight of the synapse from the pre-synaptic neuron at position *hk* to post-synaptic neuron at position *ij*^6^. 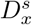 and 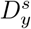 indicate the distances between the pre-synaptic neuron and the post-synaptic neurons along the horizontal and vertical axis of the unisensory area. The excitatory Gaussian function is defined by parameters 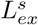 and 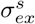, whereas the inhibitory is defined by 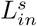 and 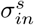. A null term (i.e. zero) was included in equation 1 to avoid auto-excitation. The change in the E/I ratio was modelled by increasing or decreasing the value of the parameter 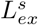 in both unisensory areas.

#### 2.4.2. Failures in top-down signalling

The failures in top-down signalling were implemented by uniformly weakening top-down synaptic weights. This was achieved by changing the value of the parameter 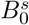, defined in equations 2 and 3:

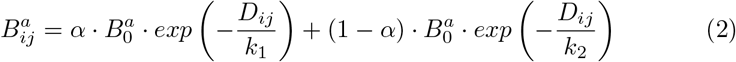

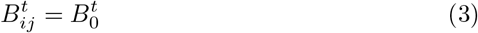

Here, the distance *D_ij_* is equal to zero for the auditory neurons that encode the first *Lim* cm of the auditory space, whilst for the neurons outside this boundary *D_ij_* is the minimum Euclidean distance between its RF centre and this boundary. 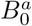 denote the value of the feedback and feedforward synapses respectively when *D_ij_* is equal to zero. *k*_1_, *k*_2_ and *α* are parameters governing the exponential decay of synaptic weights of auditory neurons encoding regions outside the near space of the hand.

#### 2.4.3. Decrease of synaptic density

The decrease of synaptic density in bottom-up and top-down connections was implemented by re-setting the connection weights of synapses that were below a certain threshold *ρ* to zero, in line with previous implementations of pruning mechanisms (Hoffman and McGlashan, 2006; Hoffman and Dobscha, 1989; Hoffman et al., 1995). We chose to prune only auditory synapses because synapses from the tactile neurons to the multisensory neuron have a uniform value and would not be affected by this implementation. Similarly, we decided to prune only auditory feedforward synapses because pruning auditory feedback synapses does not have an influence in the RT of the network (see Supplementary material for an illustration).

The extent of the pruning in the connectivity of feedforward synapses was calculated by dividing the sum of the weights of the pruned connections over the total sum of the weights of the connections before pruning.

## 3. Results

### 3.1. Parameter exploration

We systematically varied the parameters governing recurrent excitation (*L_ex_*), top-down weights 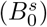 and pruning (*ρ*) to explore plausible mechanisms behind the changes in the PPS observed in SCZ and H-SPQ (see Figure 2). Our simulations revealed that an increase in both recurrent excitation and top-down synaptic weights reduces the size of the PPS representation generated by the model. In contrast, the decrease of synaptic density in auditory feedforward connections influences both the size and the slope of the PPS. Specifically, the increase of the pruning level reduces the size of the PPS and sharpens its boundary (i.e. causes the slope of the PPS representation to be steeper). However, to reproduce the size and the slope of the PPS observed in SCZ, both an increase in recurrent excitation and a decrease in synaptic density are required.

### 3.2. Identification of SCZ and H-SPQ network models

To select the models that provide the best fit to the data, we employed a simple model comparison approach based on the Root Mean Square Error (RMSE) adjusted by the number of free parameters, as defined in equation 4:

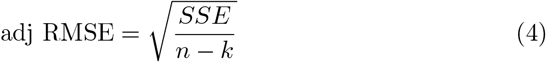

Here, SSE stands for Sum of Squared Error, n denotes the sample size and k refers to the number of free parameters.

The models that provide the best fit to the data of the SCZ and H-SPQ groups are presented in Table 1 and Figure 3. Our results suggest that the network requires increased recurrent excitation in the unisensory areas to generate PPS representations that match the data of both SCZ and H-SPQ groups. Furthermore, the network requires pruning to reproduce the PPS observed in SCZ (adj RMSE = 2.60 ms), whereas no pruning is required to match the representation observed in H-SPQ (adj RMSE = 2.22 ms).

**Table 1:** Parameters obtained by the fitting procedure. 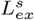 denotes the amplitude of lateral excitatory connectivity manipulated to modulate the E/I balance in the unisensory areas. Pruning (%) refers to the percentage of pruned feedforward auditory synapses. The terms a and b represent the slope and the intercept of the linear relationship.

| | $L_{ex}^s$ | Pruning (%) |
| --- | --- | --- |
| **HC** | 0.15 | 0 |
| **H-SPQ** | 1.26 | 0 |
| **SCZ** | 0.99 | 9.74 |

In this process, we discarded a model based only on recurrent excitation for SCZ (adj RMSE = 15.19 ms) and models based only on pruning for both SCZ (adj RMSE = 6.53 ms) and H-SPQ (adj RMSE = 5.47 ms). We tentatively discarded a model based on both pruning and recurrent excitation for H-SPQ (adj RMSE = 1.96 ms) because it does not provide a substantial improvement in goodness of fit compared with the recurrent excitation model (see also, Discussion).

### 3.3. Predictions

The effects of strong recurrent excitation in unisensory areas were further explored under two-point tactile stimulation, emulating the two-point discrimination task (Chang and Lenzenweger, 2001, 2005). We simulated the administration of two tactile stimuli separated by 2 cm in the modelled hand (see Supplementary Material for details in the implementation). The corresponding tactile activity in the HC and SCZ models is presented in Figure 4.

**Figure 4:**
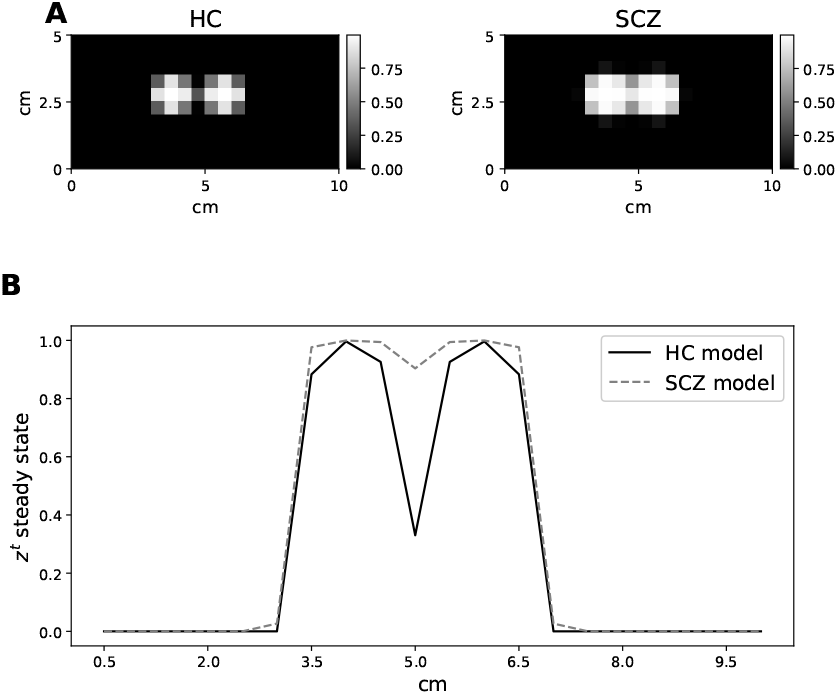
Simulation of the two-point tactile discrimination task in the HC and SCZ models. **Panel A** shows the steady-state activity of the tactile neurons of both models for two tactile stimuli presented simultaneously at positions (4cm, 2.5cm) and (6cm, 2.5cm). The grayscale represents the firing rate of the neurons that encode a given coordinate of the tactile area. The spread of activity beyond the coordinates at which the stimuli were delivered is larger in the SCZ model (right). **Panel B** shows the steady-state activity of the tactile neurons located at the 2.5cm coordinate of the y-axis in both models. The firing rates of the neurons show two clearly differentiated peaks in the HC model (solid), whereas these peaks become less distinguishable in the SCZ model (dashed).

Results of the simulation reveal that tactile activity in the SCZ model is more spread out in cortical space, leading to overlapping representations of the two stimuli. In contrast, tactile activity generated by the two stimuli does not overlap in the HC model. Using a simple read-out of activity, these differences in representations would translate into differences in discrimination performances, with higher discrimination threshold in SCZ.

## 4. Discussion

This study aimed to model the neural mechanisms that give rise to the PPS representations observed in SCZ patients and H-SPQ individuals respectively. For this purpose, an existing network model of PPS (Serino et al., 2015a) was modified in three ways: modulation of the E/I balance, weakening of top-down synapses and decreasing synaptic density between unisensory and multisensory neurons. These modifications can be thought of mimicking the consequences of the hypofunction of NMDA receptors and GABA neurons together with elevated activity of the D_2_ receptor, which are also thought to be the origin of predictive coding impairments in contemporary computational accounts of SCZ (Friston et al., 2016; Sterzer et al., 2018).

Fitting to experimental data (Section 3.2) suggests two neural mechanisms accounting for the PPS representations observed in SCZ patients and H-SPQ individuals. In the model, the PPS of H-SPQ individuals can be accounted for by an increased recurrent excitation of unisensory neurons responsible for the encoding of the spatial location of both auditory and tactile stimuli. We propose that such increase in excitation causes the narrower PPS empirically observed in H-SPQ individuals (Di Cosmo et al., 2017).

In contrast, the PPS of SCZ patients is accounted for by an increase of recurrent excitation of the unisensory neurons along with a decrease of synaptic density between unisensory and multisensory neurons. We suggest that such increase in excitation causes the small PPS observed in SCZ patients, whereas the decrease in synaptic density causes the sharply defined slope of the PPS representation registered in this population (Di Cosmo et al., 2017).

### 4.1. A novel working model of PPS representation in SCZ spectrum disorders

Our findings, as well as the behavioural data we are fitting, stand against the initially proposed working model (Noel et al., 2017) of PPS representation in SCZ, which assumed that impairments in self demarcation are related to a shallower definition of the PPS boundary originating from weaker synaptic connections between multisensory neurons and unisensory neurons. In contrast, we suggest that the reduction in the size of the PPS observed in SCZ and H-SPQ individuals is related to changes in E/I balance, leading to increased excitation of the unisensory neurons that encode the spatial location of external stimuli. We also propose that the sharper definition of the PPS boundary (i.e. the steeper slope of the PPS representation) observed in SCZ is related to a decrease in synaptic density between unisensory and multisensory neurons.

Furthermore, this new model suggests that schizotypal traits and schizophrenia differ regarding the encoding of the PPS. Although both are characterised by an E/I imbalance, we suggest that they differ in the amplitude of the decrease in synaptic density between unisensory networks and multisensory networks located in parieto-frontal areas: the decrease in H-SPQ individuals is weak or null, whereas in SCZ patients it is significant. This synaptic impairment is thought to be a consequence of failures in signalling across different levels of the cortical hierarchy produced by the E/I imbalance observed in SCZ (Friston et al., 2016).

A graphical representation of the novel working model of PPS representation in SCZ is presented in Figure 5. This figure shows the influence that the evaluated parameters (i.e. 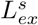 and *ρ*) have in the PPS representation, tactile discrimination and symptoms observed in SCZ patients.

**Figure 5:**
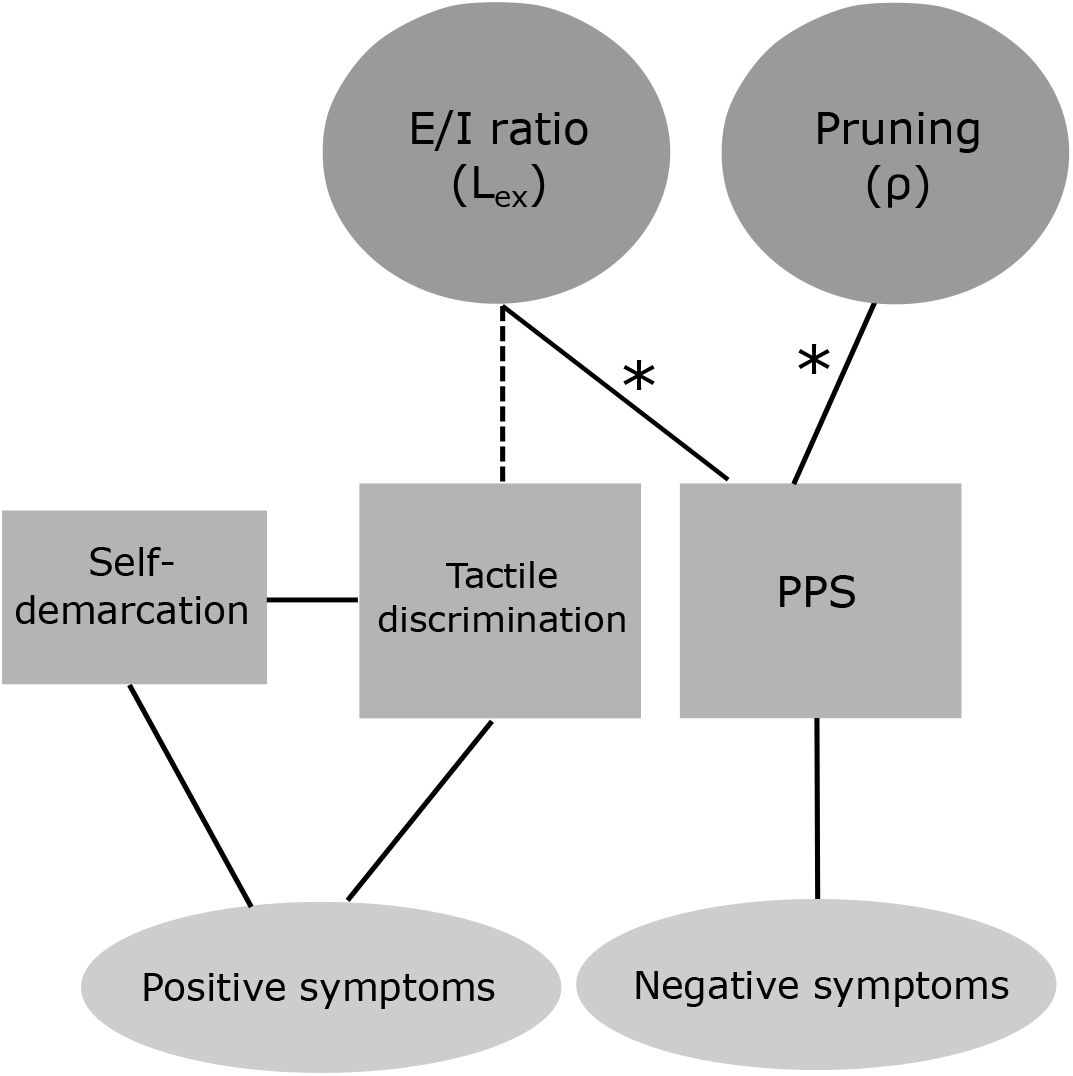
Proposed working model of PPS representation in SCZ. Solid lines marked with an asterisk represent causal relationships observed in this study; solid lines represent links reported in the literature; dashed lines represent our model’s predictions. An increased E/I ratio and pruning influence behavioural observations of tactile discrimination, PPS and their associated symptoms.

### 4.2. Encoding of PPS and tactile discrimination in SCZ

The aforementioned features of the SCZ network model are compatible with reports of impaired tactile discrimination in the SCZ spectrum. More precisely, it has been reported that SCZ patients, their relatives and H-SPQ individuals require a larger distance to detect differences in stimulation in an experimental paradigm named ‘two-point discrimination’ ^7^ (Chang and Lenzenweger, 2001, 2005; Lenzenweger, 2000; Michael and Park, 2016). In the model, such reduced tactile discrimination can be accounted for by increased recurrent excitation in unisensory neurons: tactile stimulation elicits the response of a group of neurons encoding a larger area than in control participants (due to increased E/I ratio) (see Figure 4), and the minimum distance required by the individual to detect the presence of two different stimuli increases.

Furthermore, this finding supports the view that impaired tactile sensitivity is related to self-disturbances (Ferri et al., 2016; Nelson et al., 2009; Nelson and Sass, 2017) and failures in bodily self-consciousness (Costantini et al., 2020; Noel et al., 2017). Specifically, a reduced performance in the two-point discrimination task correlates with higher scores of the cognitive-perceptual factor of the SPQ scale (Chang and Lenzenweger, 2001, 2005) and the propensity to experience the Pinocchio Illusion (PI) (Michael and Park, 2016). Similarly, reduced accuracy in the finger localisation task correlates with the strength of the Rubber Hand Illusion (RHI) and higher scores of the Schizophrenia Proneness Instrument (SPI-A) Costantini et al. (2020). In this context, we predict that disruptions of the cortical E/I balance of networks that encode portions of the skin underlie the proneness of SCZ patients to experience self-aberrations as the ones examined in the PI (Michael and Park, 2016) and the RHI (Costantini et al., 2020; Thakkar et al., 2011) paradigms.

We suggest that the increase of E/I ratio of tactile neurons influences the Bayesian sensory inference process of body ownership in the RHI (Samad et al., 2015) by increasing the likelihood that spatial and temporal signals^8^ have a single cause in the environment. We predict that such impairment weakens tactile discrimination and, as a consequence, increases the threshold at which visual and tactile stimulation are perceived as asynchronous. In turn, the increased E/I ratio can induce the illusion of owning the rubber hand, in line with the mechanics of the Bayesian model (Samad et al., 2015).

At the neurobiological level, our model is in line with reports of multisensory temporal discrimination being influenced by changes in E/I regime. Specifically, concentrations of glutamatergic compounds (Glx) conditioned on individual E/I genetic profiles account for audio-tactile temporal discrimination in the Simultaneity Judgement (SJ) task^9^ and cognitive-perceptual scores of the SPQ scale (Ferri et al., 2017): higher concentrations of Glx during the SJ task correlate with lower temporal discrimination and higher schizotypy in participants with a genetic shift towards greater excitation. In a similar magnetic resonance spectroscopy paradigm, higher E/I ratio consistently predicts higher discrimination thresholds (i.e. decreased accuracy in discrimination) in a tactile frequency discrimination task (Puts et al., 2011, 2015, 2017; Sapey-Triomphe et al., 2019). Based on this evidence, we suggest that a disruption of the cortical E/I balance is behind the decreased tactile spatial (Chang and Lenzenweger, 2001, 2005; Costantini et al., 2020; Lenzenweger, 2000) and temporal (Ferri et al., 2016) discrimination found in the SCZ spectrum and the above described symptomatology.

### 4.3. Limitations and Future Directions

The published version of the network model (Serino et al., 2015a) does not capture scaled differences in RT to the tactical stimulation when the sound is perceived at different distances from the hand^10^. This was made possible due to the additional use of a linear regression matching the output of the network and behavioural data. Hence, the network model accurately reproduces the shape of the PPS representation (i.e. central point and slope), but it is not able to reproduce realistic RT changes across distance points.

The experimental evidence directly assessing the PPS representation in the SCZ spectrum has only started to accumulate (Di Cosmo et al., 2017; Ferroni et al., 2020). Our findings are based on the modelling of a single study that employs a robust experimental paradigm to behaviourally measure the PPS (Canzoneri et al., 2012). To further progress in this area, further studies with more recent behavioural (Hobeika et al., 2020; Serino et al., 2015b) and neurocognitive measures (Naro et al., 2019; Noel et al., 2019a, b) of the PPS representation in the SCZ spectrum are needed.

Our model provides a starting point to identify the neural mechanisms behind the changes observed in the PPS representation in SCZ. Further studies should be aimed at clarifying the specific mechanism behind the proposed E/I imbalance since our model is not able to distinguish whether it is produced by increased excitation or decreased inhibition ^11^. Similarly, further research should explore the specific neural mechanism behind the proposed dysconnection between multisensory and unisensory networks sincer our approach is not able to assess the effects of pruning feedback synapses ^12^. Further anatomical and neurophysiological studies will be needed to answer these questions.

Finally, further computational studies should aim at exploring changes in the PPS representation during learning tasks (e.g. Ferroni et al. (2020); Noel et al. (2018b)). This would open up the possibility to explore Bayesian predictive coding computations (i.e. predictions, prediction errors and precision estimations), which are central in contemporary accounts of SCZ (Adams et al., 2013; Friston et al., 2016; Sterzer et al., 2018). In addition, further computational studies should aim at elucidating the neural mechanisms behind impairments in tactile (Ferri et al., 2016) and audio-tactile (Di Cosmo et al., unpublished; Ferri et al., 2017) temporal discrimination found in the SCZ spectrum. This could be complemented with the study of the integration of visual-somatosensory temporal and spatial signals involved in the strong RHI observed in SCZ (Costantini et al., 2020; Thakkar et al., 2011). Overall, further work could attempt to relate such a neural network implementation and the predictive coding framework whose neural basis has still to be fully established Shipp (2016).

## Footnotes

1 These experiments were built to manipulate feelings of body ownership in participants as a result of controlled multimodal stimulation.

2 The RHI is an experimental paradigm designed to generate in participants the feeling that a rubber hand is their own, as a result of synchronous or asynchronous stimulation of the fake hand and their real hand (which is not visible to participants).

3 The PI is an experimental paradigm designed to generate in participants the sensation that their nose is growing in response to tactile-proprioceptive stimulation.

4 The code employed for the implementation of the model can be found in: https://github.com/renatoparedes/SCZ_PPS_model

5 Although the exact value of the velocity of sound was not reported in the aforementioned study (Di Cosmo et al., 2017), further studies employing the audio-tactile paradigm have reported velocities ranging from 22 cm/s to 35 cm/s (Serino et al., 2015b)

6 Here, *i* and *j* denote the position of the neuron in the unisensory area *s*, where *s* can be either *t* (tactile) or *a* (auditory). Refer to Supplementary material for a detailed description.

7 In this paradigm, two-point stimuli are administered by an aesthesiometer in the palm of the hand at different distances to detect the threshold at which an individual stops perceiving two separated stimuli.

8 Bayesian modelling of the RHI (Samad et al., 2015) proposed that spatial signals are composed by visual and proprioceptive information of the hand location, whereas temporal signals are composed by the synchronisation of visual and tactile stimulation

9 In the SJ task participants had to indicate whether auditory and tactile stimuli delivered at changing onset asynchronies were presented together or not.

10 For example, the network of the SCZ model registers differences of less than 5 ms across the distance points, whereas differences of approximately 20 ms are observed in the experimental data of patients with SCZ.

11 With a different parameterisation, our model is able to reproduce changes in the size of the PPS by diminishing the value of 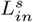 instead of increasing 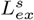.

12 Our pruning implementation only influences RT when impairing auditory feedforward synapses because of the specific setup of top-down and bottom-up synapses defined in the published version of the network model (see Supplementary materials for details).

## Notes

https://github.com/renatoparedes/SCZ_PPS_Model

### Competing Interest Statement

The authors have declared no competing interest.

