## Supplementary Material for "Influence of E/I balance and pruning in peri-personal space differences in schizophrenia: a computational approach"

November 6, 2020

#### 1 Neural network model

##### 1.1 Network architecture

The model is composed of two recurrently connected unisensory areas (auditory and tactile) interacting with one multisensory area (as shown in Figure 1). Each neuron in both unisensory areas has its own receptive field (RF) defined in hand-centred in coordinates (along the horizontal and vertical axis) and responds to stimulation received by an external input with specific spatial coordinates in relation to the hand. In the following, $x_i^s$ and $y_j^s$ will denote the centre of the RF of a given neuron at position $ij$ of the unisensory area $s$, where $s$ can be either $t$ (tactile) or $a$ (auditory).

The tactile area is composed of 200 neurons (disposed in a $M^t = 20$ x $N^t = 10$ grid) that encode a skin portion of 10 cm x 5 cm corresponding to the left hand of an individual. This area may roughly represent high-order somatosensory areas in the parietal lobe [1, 2]. The tactile RF centres coordinates are defined by Equation 1.

$$x_i^t = i \cdot 0.5cm(i = 1, 2, .., M^t) \qquad y_j^t = j \cdot 0.5cm(j = 1, 2, .., N^t) \tag{1}$$

Similarly, the auditory area is composed of 60 neurons (disposed in a $M^a = 20$ x $N^a = 3$ grid) that cover an auditory space of 200 cm x 30 cm on and around the hand. Here it must be noted that the auditory area is composed of fewer neurons that encode larger spaces to mimic the lower spatial resolution of the auditory system. Hence, this auditory area roughly resembles high-order neural structures of the auditory pathway that encode the spatial location of the sound source [3]. The auditory RF centres coordinates are defined by Equation 2.

$$x_i^a = i \cdot 10 - 5cm(i = 1, 2, .., M^a) \qquad y_j^a = j \cdot 10 - 15cm(j = 1, 2, .., N^a) \tag{2}$$

Finally, the multisensory area is composed of a single neuron that is connected to all neurons in both auditory and tactile areas. This design is congruent with the observation that only few multisensory neurons are capable of encoding the entire space of the hand [4, 5, 6, 7].

#### 1.2 Network synapses

Each neuron of the unisensory areas is recurrently connected to all neurons in the area to which it belongs. These synaptic connections are symmetrical and are arranged in a "Mexican hat" pattern, which promotes excitation of neurons that are closer and fosters inhibition in neurons that are further. In this setup, an external stimulus activates a limited number of unisensory neurons and avoids the propagation of excitation in the unisensory area. The specific weights assigned to the synapses are obtained as the difference between two Gaussian functions (one excitatory and one inhibitory), according to equation 3.

$$L^s_{ij,hk} = \begin{cases} L^s_{ex} \cdot exp\left(-\frac{(D^s_x)^2+\left(D^s_y\right)^2}{2\cdot(\sigma^s_{ex})^2}\right) - L^s_{in} \cdot exp\left(-\frac{(D^s_x)^2+\left(D^s_y\right)^2}{2\cdot\left(\sigma^s_{in}\right)^2}\right), & ij \neq hk \\ 0, & ij = hk \end{cases} \tag{3}$$

$$s = t, a$$

$L^s_{ij,hk}$ denotes the weight of the synapse from the pre-synaptic neuron at position *hk* to post-synaptic neuron at position *ij*. $D^s_x$ and $D^s_y$ indicate the distances between the pre-synaptic neuron and the post-synaptic neurons along the horizontal and vertical axis of the unisensory area. The excitatory Gaussian function is defined by parameters $L^s_{ex}$ and $\sigma^s_{ex}$, whereas the inhibitory is defined by $L^s_{in}$ and $\sigma^s_{in}$. A null term (i.e. zero) is included in equation 3 to avoid auto-excitation.

The synaptic strength between the multisensory neuron and the unisensory neurons are different according to each modality. The weights of the synapses that connect the tactile neurons and the multisensory neuron are all the same independent of the position of the neuron in the tactile area, according to Equations 4 and 5. $W^t_{ij}$ and $B^t_{ij}$ denote the weight of the feedforward and feeedback tactile synapses respectively. In turn, $W^t_0$ and $B^t_0$ represent the fixed value assigned to those synapses.

$$W^t_{ij} = W^t_0 \tag{4}$$

$$B^t_{ij} = B^t_0 \tag{5}$$

In contrast, the weights of the synapses that connect the auditory neurons and the multisensory neuron depend on the portion of auditory space that each neuron encodes with respect to the modelled skin portion of the hand. Thus, the synapses hold a constant weight for the space on and near the hand, covering

$Lim$ cm of the auditory space (20 cm for the hand and $Lim$-20 cm for the space close to the hand). The synaptic weights outside this boundary (i.e. $Lim$) are described by a bi-exponential function decreasing with the distance between the neurons' RF and the hand, as shown by Equations 7 and 6. $W_{ij}^a$ and $B_{ij}^a$ denote the weight of the feedforward and feedback auditory synapses respectively.

$$W_{ij}^a = \alpha \cdot W_0^a \cdot exp\left(-\frac{D_{ij}}{k_1}\right) + (1-\alpha) \cdot W_0^a \cdot exp\left(-\frac{D_{ij}}{k_2}\right) \tag{6}$$

$$B_{ij}^a = \alpha \cdot B_0^a \cdot exp\left(-\frac{D_{ij}}{k_1}\right) + (1-\alpha) \cdot B_0^a \cdot exp\left(-\frac{D_{ij}}{k_2}\right) \tag{7}$$

In both equations, the distance $D_{ij}$ is equal to zero for the auditory neurons that encode the first $Lim$ cm of the auditory space, whilst for the neurons outside this boundary $D_{ij}$ is the minimum Euclidean distance between its RF centre and this boundary. $W_0^a$ and $B_0^a$ denote the value of the feedback and feedforward synapses respectively when $D_{ij}$ is equal to zero. $k_1$, $k_2$ and $\alpha$ are parameters governing the exponential decay of synaptic weights of auditory neurons encoding regions outside the near space of the hand. The values assigned to these later parameters were selected to produce a fast decay immediately outside this space and a low decay afterwards, as in the original publication [8]. The feedforward and feedback auditory synaptic weights are shown in Figure 1.

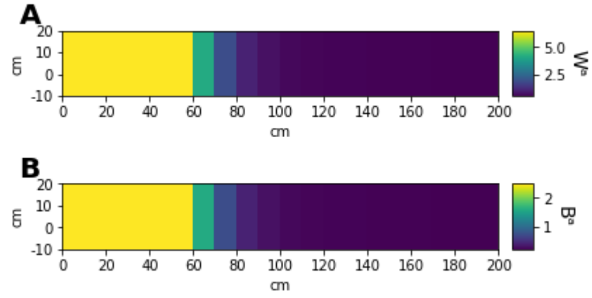


Figure 1: Bottom-up and top-down connections between the auditory area and the multisensory area. **Panels A** and **B** show feedforward ($W^a$) and feedback ($B^a$) auditory synapses respectively. The colours represent the strength of the synaptic connections into neurons encoding the auditory area. The images illustrate stronger synapses towards neurons encoding the space close to the hand and exponentially weaker ones outside this region.

#### 1.3 Network activity

The model is composed of non-spiking (rate) neurons, whose output is a continuous variable representing the neuron's firing rate. Each neuron of the model responds to its overall input through first-order temporal dynamics and a sigmoidal transfer function. The unisensory neurons temporal dynamics are defined in equation 8:

$$\tau \frac{dq_{ij}^{s}\left(t\right)}{dt} = -q_{ij}^{s}\left(t\right) + u_{ij}^{s}\left(t\right), \qquad s = t, a \tag{8}$$

In equation 8, $q_{ij}^{s}\left(t\right)$ stands for the state variable of a neuron at a given time step. This is affected by $u_{ij}^{s}\left(t\right)$, which denotes the overall input of a neuron at a given time step. Briefly, this input is the sum of the external stimulation received convolved with its RF, the recurrent input received from other neurons of its same area and the feedback input of the multisensory area. In this temporal relationship, $\tau$ is the time constant of the differential equation and its value is set according to the original publication [8], in agreement with realistic membrane time constants reported in the literature [9].

The unisensory neurons input $u_{ij}^{s}\left(t\right)$ is computed by Equation 9.

$$u_{ij}^{s}\left(t\right) = \varphi_{ij}^{s} + I_{ij}^{s}\left(t\right) + b_{ij}^{s}\left(t\right), \qquad s = t, a \tag{9}$$

The term $\varphi_{ij}^{s}$ denotes the input due to external stimulus, $I_{ij}^{s}\left(t\right)$ indicates the input received from other neurons in the same area and $b_{ij}^{s}\left(t\right)$ represents the feedback input from the multisensory neuron.

The input $\varphi_{ij}^{s}$ is defined by Equation 10.

$$\varphi_{ij}^{s} = \sum_{l}\sum_{n} \Phi_{ij}^{s}(x_l, y_n) \cdot I^{s}\left(x_l, y_n\right) \Delta x_l \Delta y_n, \qquad s = t, a \tag{10}$$

The expression $I^{s}\left(x_l, y_n\right)$ denotes the external stimulus applied at coordinates x and y, as defined in Equation 20. The term $\Phi_{ij}^{s}(x_l, y_n)$ represents the RF of the unisensory neuron $ij$ in the unisensory area $s$. Equation 10 is solved by considering $\Delta x_l = \Delta y_n = 0.2cm$.

The RF $\Phi_{ij}^{s}(x_l, y_n)$ is defined by the Gaussian function presented in Equation 11.

$$\Phi_{ij}^{s}(x, y) = \Phi_0^{s} \cdot exp\left(-\frac{\left(x_i^{s} - x\right)^2 + \left(y_j^{s} - y\right)^2}{2 \cdot \left(\sigma_{\Phi}^{s}\right)^2}\right), \qquad s = t, a \tag{11}$$

The terms $\mathrm{x_i}$ and $\mathrm{y_j}$ denote the coordinates of the RF centre, whereas x and y represent spatial coordinates within the unisensory areas. The parameters $\Phi_0^{s}$ and $\sigma_{\Phi}^{s}$ govern the amplitude and standard deviation of the Gaussian function that define the RF of the unisensory neurons.

Following with Equation 9 description, the lateral inputs $I_{ij}^{s}\left(t\right)$ were calculated by Equation 12.

$$I_{ij}^{s}(t) = \sum_{h=1}^{N^s} \sum_{k=1}^{M^s} L_{ij,hk}^{s} \cdot z_{hk}^{s}(t), \qquad s = t, a \tag{12}$$

The term $L_{ij,hk}^{s}$ indicates the synaptic weights calculated by Equation 3. Similarly, $z_{hk}^{s}(t)$ denotes the neuron activation at time $t$, as described by equation 14.

Finally, the feedback input $b_{ij}^{s}(t)$ was computed by Equation 13.

$$b_{ij}^{s}(t) = B_{ij}^{s} \cdot z^{m}(t), \qquad s = t, a \tag{13}$$

The term $B_{ij}^{s}$ represents the strenght of feedback synapses, as described by equation 7. The expression $z^{m}(t)$ denotes the activation state of the multisensory neuron at time $t$.

The unisensory neurons activity $z_{ij}^{s}$ is computed out of the state variable $q_{ij}^{s}(t)$, as shown in equation 14:

$$z_{ij}^{s}(t) = \Psi\left(q_{ij}^{s}(t)\right) \cdot H\left(\Psi\left(q_{ij}^{s}(t)\right)\right), \qquad s = t, a \tag{14}$$

Equation 14 displays a sigmoidal function $\Psi$ being applied to the state variable $q_{ij}^{s}(t)$. The result of this is multiplied by a Heaviside function $H$ to avoid negative values in the activity of neurons. Finally, the sigmoidal function $\Psi$ is defined by equation 15:

$$\Psi\left(q_{ij}^{s}(t)\right) = \frac{f_{min}^{s} + f_{max}^{s} \cdot e^{\left(q_{ij}^{s} - q_{c}^{s}\right) \cdot r^{s}}}{1 + e^{\left(q_{ij}^{s} - q_{c}^{s}\right) \cdot r^{s}}}, \qquad s = t, a \tag{15}$$

Here, $f_{min}^{s}$ and $f_{max}^{s}$ are the lower and upper boundaries of the sigmoid function. $q_{c}^{s}$ is the central point of the sigmoid and $r^{s}$ denotes the slope of the curve at the central point.

The multisensory neuron activity is defined by an analogous set of equations:

$$\tau \frac{dq^{m}(t)}{dt} = -q^{m}(t) + u^{m}(t) \tag{16}$$

$$z^{m}(t) = \Psi\left(q^{m}(t)\right) \cdot H\left(\Psi\left(q^{m}(t)\right)\right) \tag{17}$$

$$\Psi\left(q^{m}(t)\right) = \frac{f_{min}^{m} + f_{max}^{m} \cdot e^{\left(q^{m} - q_{c}\right) \cdot r^{m}}}{1 + e^{\left(q^{m} - q_{c}\right) \cdot r^{m}}} \tag{18}$$

The term $u^{m}(t)$ represents the multisensory neuron input described by Equation 19. The description of the remaining terms are analogous to Equations 8,14 and 15.

The input $u^m(t)$ received by the neuron is composed of the sum of the feed-forward inputs from both auditory and tactile areas, as presented in equation 19.

$$u^m(t) = \sum_{i=1}^{N^t} \sum_{j=1}^{M^t} W_{ij}^t \cdot z_{ij}^t(t) + \sum_{i=1}^{N^a} \sum_{j=1}^{M^a} W_{ij}^a \cdot z_{ij}^a(t) \tag{19}$$

Here, 19, $z_{ij}^s(t)$ (s = t,a) denotes the activity of neuron $ij$ in its respective unisensory area $s$. This is obtained by equations 14, 15 and 19. In addition, $W_{ij}^s$ (s = t,a) represents the feedforward synapses from the unisensory neuron $ij$ in the area $s$ to the multisensory neuron. These synapses were introduced in the previous subsection and displayed in Figure 1A.

The reaction time (RT) of the network was registered at each distance point as the time at which any neuron of the tactile area reached the 90% of its maximum activation state (i.e. 90% of $f_{max}^t$).

A couple of remarks must be considered before concluding this subsection. First, all the aforementioned differential equations were solved numerically employing the Euler integration method with a discrete time step of 0.4 ms, as in the original implementation [8]. Second, as highlighted by the authors of the original model [10, 11], each neuron of this network must not be interpreted as a single cell only but as the average behaviour of a group of cells that approximately share the same RF and code for a similar stimulus location. A similar interpretation must be held for the synapses among the neurons of the network: they do not represent a single synaptic connection but a summary of the overall synaptic strength of a group of neurons.

#### 1.4 Network stimuli

Network stimuli was built for both unisensory areas to resemble the stimulation received by participants in the experiment described in Section 2.1. Both auditory and tactile stimuli had their centres located at a specific coordinate of their respective unisensory area. These inputs were implemented by a bidimensional Gaussian function $I^s$, as defined in equation 20. In contrast to the original implementation [8], we opted to exclude the random noise introduced in the stimulus intensity to avoid unexplained network variability [1].

$$I^s(x,y) = I_0^s \cdot exp\left(-\frac{(x_0^s - x)^2 + (y_0^s - y)^2}{2 \cdot (\sigma_I^s)^2}\right), \qquad s = t,a \tag{20}$$

Here, $I_0^s$ denotes the intensity of the stimulus. $x_0^s$ and $y_0^s$ are the coordinates of the central point of the stimulus applied in the unisensory area $s$. $\sigma_I^s$ represents the spatial of the stimulus and was given a small value to simulate localised stimuli.

[1] This change causes the model to be fully deterministic and be influenced only by the previously described network mechanics.

The tactile stimuli was always applied at coordinate $x^t = 5$ cm and $y^t = 2.5$ cm. In contrast, the auditory stimuli was always applied at coordinate $y^a = 5$ cm but the $x^a$ coordinate changed across trials to simulate the presentation of the sound stimuli at different distances from the hand.

#### 1.5 Sigmoid fitting of network output

The RTs values generated by the network in the experiment simulation were fitted to a sigmoid function described by Equation 21.

$$f_{RT}(D) = \frac{\eta_{min} + \eta_{max} \cdot e^{\frac{(D-D_c)}{h}}}{1 + e^{\frac{(D-D_c)}{h}}} \tag{21}$$

The term $D$ denotes the distance from the hand in cm at which the auditory stimulus was applied. The parameters $\eta_{min}$ and $\eta_{max}$ represent the lower and upper saturation of the sigmoidal relationship. $D_c$ is the central point of the function and $h$ denotes the slope of the function at its central point.

Parameters $D_c$ and $h$ were estimated by a least-squares fitting procedure using the Trust Region Reflective algorithm available in the SciPy v.1.4.1 library for the Python programming language [12]. The central point was bounded to be positive, whereas the slope was unbounded. The initial guesses given to the algorithm were defined according to equations 22 and 23.

$$D_{c_{in}} = \frac{max(D)}{2} \tag{22}$$

$$h_{in} = \frac{max(RT) - min(RT)}{4 \cdot \left(\frac{max(RT)-min(RT)}{max(D)-min(D)}\right)} \tag{23}$$

Here, $D_{c_{in}}$ and $h_{in}$ denote the initial guesses of the central point and the slope respectively. The arguments *max* and *min* refer to the maximum and minimum value of a given vector respectively. The estimated value of the sigmoid central point ($D_c$) was assumed as the boundary of the PPS representation generated by the model and the value of the slope ($h$) the sharpness of its definition.

#### 2 Values of HC model parameters

| | | | |
|---|---|---|---|
| Unisensory receptive fields | | | |
| $\phi_0^t = 1$ | $\sigma_\phi^t = 0.5$ | $\phi_0^a = 1$ | $\sigma_\phi^a = 10$ |
| External stimuli | | | |
| $I_0^t = 2.5$ | $\sigma_I^t = 0.3$ cm | $I_0^a = 3.6$ | $\sigma_I^a = 0.3$ cm |
| Lateral synapses in unisensory areas | | | |
| $L_{ex}^t = 0.15$ | $L_{in}^t = 0.05$ | $\sigma_{ex}^t = 1$ cm | $\sigma_{in}^t = 4$ cm |
| $L_{ex}^a = 0.15$ | $L_{in}^a = 0.05$ | $\sigma_{ex}^a = 20$ cm | $\sigma_{in}^a = 80$ cm |
| Feedforward and feedback synapses | | | |
| $W_0^t = 6.5$ | $W_0^a = 6.5$ | $B_0^t = 2.5$ | $B_0^a = 2.5$ |
| $k_1 = 16.71$ cm | $k_2 = 813.75$ cm | $\alpha = 0.98$ | Lim = 48.68 cm |
| Input-output relationship of unisensory neurons | | | |
| $f_{mim}^t$ = -0.12 | $f_{max}^t = 1$ | $q_c^t = 19.43$ | $r^t = 0.34$ |
| $f_{mim}^a$ = -0.12 | $f_{max}^a = 1$ | $q_c^a = 19.43$ | $r^a = 0.34$ |
| $\tau = 20$ ms | | | |
| Input-output relationship of the multisensory neuron | | | |
| $f_{mim}^m = 0$ | $f_{max}^m = 1$ | $q_c^m = 12$ | $r^m = 0.6$ |
| $\tau = 20$ ms | | | |

Table 1: Values of the parameters in the HC model. This parameterisation remained fixed throughout the simulations computed to identify the SCZ and H-SPQ models.

### 3 Modelling the influence of SCZ and H-SPQ in the network

#### 3.1 Decrease of synaptic density

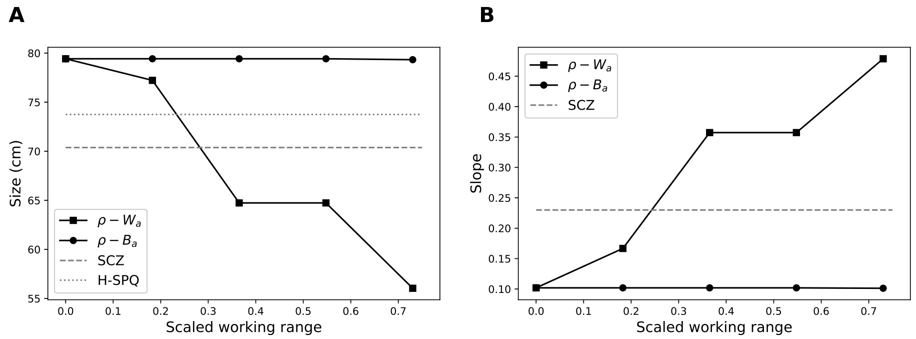


Figure 2: Effects of pruning auditory synapses in the size and the slope of the PPS representation. **Panels A** and **B** show the effects of systematic variation of the $\rho$ parameter applied to feedforward ($W_a$) and feedback ($B_a$) synapses in the range at which it produces sigmoid-like PPS representations. The dashed lines indicate the size and slope reported in the experimental study [13]. The changes in the size and the slope of the PPS observed in SCZ and H-SPQ could be reproduced by pruning of auditory feedforward synapses. In contrast, pruning of auditory feedback synapses does not have an effect on the PPS representation.

### 4 Fitting procedure

A fitting procedure inspired by an established method for parametrisation of connectionist models [14] was employed [2]. This procedure consisted in matching the time units of the model and the experiment and minimising a cost function with an stochastic optimisation algorithm. At every iteration of the minimisation routine, the RT were matched by a standard linear regression, as presented in Equation 24.

$$RT^{data} = a \cdot RT^{model} + b \tag{24}$$

Here, $RT^{data}$ and $RT^{model}$ represent the RTs obtained in the empirical study and the experiment simulation respectively. Parameter $a$ denotes how many milliseconds correspond to one time unit of the network model, whereas parameter $b$ represents the duration of neural processing not captured by the model.

During the optimisation, negative values of $a$ and $b$ were converted to 0 because negative coefficients do not make sense according to this definition. The employed cost function is defined by Equation 25.

$$Cost = \sum_{i=1}^{N} \left( \frac{RT_i^{data} - RT_i^{model}}{RT_i^{data}} \right)^2 \tag{25}$$

Here, $RT_i^{data}$ and $RT_i^{model}$ denote the RT measured at the $i$th distance point. N represents the number of distances measured (e.g. 5 in the empirical study).

This cost function was minimised by a fast implementation of the differential evolution algorithm [15] available in the SciPy v1.4.1 library for the Python programming language [12].

For the fitting we considered 7 equally spaced distance points in the range between 39 and 111 cm. Hence, $RT^{model}$ is composed of 7 RTs generated by the model at those distance points. Similarly, $RT^{data}$ is composed of 7 RT taken from the mean sigmoid fit obtained out of the experimental data of a given group.

#### 4.1 HC Fitting

We fitted the model to the healthy control data. Here, $k_1$, $k_2$, $\alpha$ and $Lim$ (see Equations 6 and 7) varied freely to represent individual differences in the PPS representation due to stable anatomical factors (i.e. immutable across the duration of the experiment). All the other parameters (see Table 1) remained fixed and set to the values published by [8]. The bounds given to the optimisation algorithm to find $k_1$, $k_2$, $\alpha$ and $Lim$ were (1, 50), (500, 1000), (0.25, 1) and (20, 80) respectively.

[2] The main difference between the original method and our procedure is that we omitted the steps aimed to deal with response variability because our network generates deterministic responses.

The results of the fitting to the data of healthy controls are presented in Figure 3 and Table 2. The plot reveals that the model is able to closely reproduce the control data for the non-social condition (RMSE = 0.37).

| **Model** | $k_1$ | $k_2$ | $\alpha$ | $Lim$ | $\boldsymbol{a}$ | $\boldsymbol{b}$ |
|---|---|---|---|---|---|---|
| HC | 16.71 | 813.75 | 0.98 | 48.68 | 2.73 | 137.63 |

Table 2: Parameters obtained after fitting the model to healthy control data.

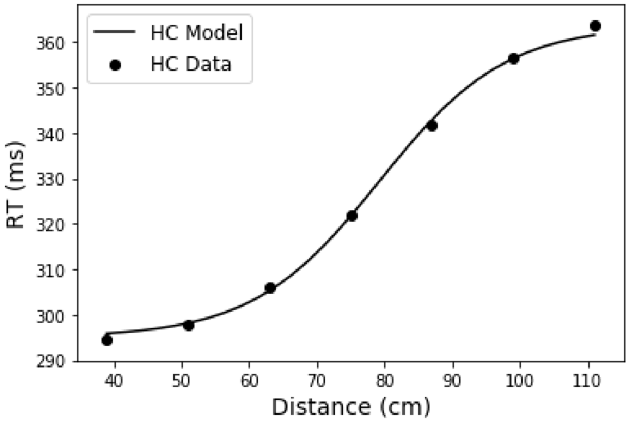


Figure 3: Results of fitting the model to the healthy control data. The line depicts the sigmoid fit obtained out of the data generated by the HC model. The dots represent samples of the sigmoid fit obtained out of the data collected in the behavioural study for such group. The quantitative fitting of the model using $k_1$, $k_2$, $\alpha$ and $Lim$ leads to a close match to the experimental data of healthy controls.

We decided to fit the parameters $a$ and $b$ in further fittings because the parameters that define the E/I balance in the model (i.e. $L^s_{ex}$), feedback input (i.e. $B^s_0$) and synaptic density decrease (i.e. $\rho$) have an effect on the scale of $RT^{model}$. Hence, the matching of time units of the model and the experiment needs to be performed every time these parameters are modified.

#### 4.2 H-SPQ Fitting

We fitted the HC model to the H-SPQ data. The bounds given to the optimisation algorithm to find $L^s_{ex}$ and $\rho$ were (0, 3) and (0, 6.5) respectively. These bounds were chosen because the PPS representation generated by values outside these range ceases to resemble a sigmoid curve, which is characteristic of the average PPS representations found in the literature for HC, SCZ and H-SPQ [13]. The results of the fitting to the data of H-SPQ individuals are presented in Table 3.

| Model | $L_{ex}^s$ | $\rho$ | $a$ | $b$ | adj RMSE |
|---|---|---|---|---|---|
| H-SPQ $L_{ex}^s$ | 1.26 | 0 | 5.11 | 1.10 | 2.22 |
| H-SPQ $\rho$ | 0.15 | 2.01 | 2.15 | 146.91 | 5.47 |
| H-SPQ $L_{ex}^s$ - $\rho$ | 1.59 | 0.57 | 5.25 | 0.18 | 1.96 |

Table 3: Parameters obtained after fitting the HC model to H-SPQ data.

#### 4.3 SCZ Fitting

We fitted the HC model to the SCZ data. As in the previous section, the bounds given to the optimisation algorithm to find $L_{ex}^s$ and $\rho$ were (0, 3) and (0, 6.5) respectively. The results of the fitting to the data of SCZ patients are presented in Table 4.

| Model | $L_{ex}^s$ | $\rho$ | $a$ | $b$ | adj RMSE |
|---|---|---|---|---|---|
| SCZ $L_{ex}^s$ | 0.82 | 0 | 6.36 | 3.35 | 15.19 |
| SCZ $\rho$ | 0.15 | 2.20 | 3.35 | 153.67 | 6.53 |
| SCZ $L_{ex}^s$ - $\rho$ | 0.99 | 2.00 | 5.61 | 43.36 | 2.60 |

Table 4: Parameters obtained after fitting the HC model to SCZ data.

### 5 Predictions

To emulate the two-point tactile discrimination task, we modified equation 20 to be able to generate two stimuli located at different coordinates of the tactile area, as described in equation 26.

$$
\begin{aligned}
I^t(x,y) = I_0^t \cdot exp\left(-\frac{\left(x_{0_a}^t - x\right)^2 + \left(y_{0_a}^t - y\right)^2}{2 \cdot \left(\sigma_I^t\right)^2}\right) \\
+ I_0^t \cdot exp\left(-\frac{\left(x_{0_b}^t - x\right)^2 + \left(y_{0_b}^t - y\right)^2}{2 \cdot \left(\sigma_I^t\right)^2}\right)
\end{aligned}
\tag{26}
$$

Here, $(x_{0_a}^t,\, y_{0_a}^t)$ and $(x_{0_b}^t,\, y_{0_b}^t)$ are the two set of coordinates of the central points of the stimuli delivered in the unisensory area $s$. The remaining terms descriptions are the same as in equation 20.

We simulated the experiment in the network model without providing auditory stimulation (i.e. $I_0^a = 0$). Hence, we increased the intensity of tactile stimulation $(I_0^t)$ from 2.5 to 3.25 to compensate for the lack of multisensory input generated by auditory stimulation.
